# Probing machine-learning classifiers using noise, bubbles, and reverse correlation

**DOI:** 10.1101/2020.06.22.165688

**Authors:** Etienne Thoret, Thomas Andrillon, Damien Léger, Daniel Pressnitzer

## Abstract

**Background:** Many scientific fields now use machine-learning tools to assist with complex classification tasks. In neuroscience, automatic classifiers may be useful to diagnose medical images, monitor electrophysiological signals, or decode perceptual and cognitive states from neural signals. However, such tools often remain black-boxes: they lack interpretability. A lack of interpretability has obvious ethical implications for clinical applications, but it also limits the usefulness of these tools to formulate new theoretical hypotheses.

**New method:** We propose a simple and versatile method to help characterize the information used by a classifier to perform its task. Specifically, noisy versions of training samples or, when the training set is unavailable, custom-generated noisy samples, are fed to the classifier. Multiplicative noise, so-called “bubbles”, or additive noise are applied to the input representation. Reverse correlation techniques are then adapted to extract either the discriminative information, defined as the parts of the input dataset that have the most weight in the classification decision, and represented information, which correspond to the input features most representative of each category.

**Results:** The method is illustrated for the classification of written numbers by a convolutional deep neural network; for the classification of speech versus music by a support vector machine; and for the classification of sleep stages from neurophysiological recordings by a random forest classifier. In all cases, the features extracted are readily interpretable.

**Comparison with Existing Methods:** Quantitative comparisons show that the present method can match state-of-the art interpretation methods for convolutional neural networks. Moreover, our method uses an intuitive and well-established framework in neuroscience, reverse correlation. It is also generic: it can be applied to any kind of classifier and any kind of input data.

**Conclusions:** We suggest that the method could provide an intuitive and versatile interface between neuroscientists and machine-learning tools.

**Highlights:**

- The heuristics of black-box classifiers can be probed with noisy inputs
- The relevant features can be visualised in the input representation space
- The method applies to any kind of data such as 2D images or 1D time series
- It applies to any classifier such as deep neural networks, support vector machines, random forests

## Introduction

Applications of machine-learning techniques permeate more and more scientific fields, with rapid and sometimes unexpected success (LeCun et al., 2015; Jordan & Mitschell, 2015; Kriegeskorte & Douglas, 2018; Richards et al., 2019). At the same time, it is becoming a widely-acknowledged issue that many of these tools are often used as black boxes, even though they would benefit from being interpreted (Molnar, 2020; Doshi-Velez & Kim, 2017). For instance, if a Deep Neural Network (DNN) was used to make life-changing decisions such as deciding on an intervention based on medical imagery, both the clinicians and the patients would have a clear desire to know the rationale that motivated the decision. Also, the power of classifiers to detect useful information in large datasets holds many promises to improve theoretical models. but then, understanding at least to some extent the classifier’s operation is crucial (Zihni et al., 2020).

Understanding what a complex classifier does after being trained on possibly millions or billions of samples is usually hard. It is hard for a reason: if the task that the classifier solves had a known explicit solution, then there probably would not have been any incentive to develop the classifier in the first place. In addition, modern techniques involve artificial network architectures with interconnected layers, each including highly non-linear operations (Sejnowski, Kienker, & Hinton, 1986). A lot of the computational power of such algorithms lies in cascades of feed-forward and feed-back non-linear operations. Unfortunately, human reasoning seems most at ease to generate intuitions with linear processes, and not for complex combinations of non-linear ones.

As a consequence, designing methods to interpret machine learning tools is a fast-growing field of research of its own right, often designated under the term *Explainable AI* (Guidotti et al., 2018). It has dedicated journals within the machine learning community (e.g. *Distill*) and an associated DARPA challenge (*XAI*). Recent reviews covering the types of methods exist (Molnar, 2020), also covering more specifically the feature visualization approach taken here (Olah et al., 2017). Within this context, our aim is not to outperform the state-of-the art specialized interpretability methods, but rather to provide a general tool that will hopefully be intuitive to neuroscientists, as it is based on familiar methods for this community. The manuscript describes the method, provides an open software library to use it, and shows examples of application, demonstrating how it can achieve useful results.

The gist of the method is to try and reveal the input features used by an automatic classifier to achieve its task *without any knowledge* about the classifier. As such, it is what is termed an “agnostic” method of explanation: it does not attempt to describe mechanistically the operation of a specific classifier, which it considers as a black box. Even if the classifier’s details are available, they are not considered as they may be too complex to understand intuitively. Rather, the aim is to relate features of the input space to the classifier’s decisions, that is to identify the features that participate the most to the classifier’s performance. Such a problem is closely related to issues that neuroscientists and experimental psychologists have been addressing for years: providing useful insights of, for instance, human perception, without a full knowledge of the highly complex and non-linear underlying processes implemented by the brain.

The method we propose is directly inspired from the reverse correlation techniques developed for studying human audition (Ahumada et al., 1971) and vision (Neri et al., 1999; Gosselin & Shyns, 2001, 2003). Reverse correlation is based on linear systems analysis (Wiener, 1966). It uses stochastic perturbations of a system to observe its output. If the system were linear, an average of the inputs weighed by the observed outputs would be able to fully characterize the system. However, even for the highly non-linear systems studied by neuroscience, reverse correlation has a track-record of successful applications. For neurophysiology, averaging input stimuli according to neural firing rates has been used to describe neural selectivity (Ringach & Shapeley, 2004 for a review). For psychophysics, averaging input stimuli according to participant’s decisions has revealed stimulus features on which such decisions are made for detection or discrimination tasks (Ahumada et al., 1971; Gosselin & Shyns, 2001, 2002). In this spirit, it seems appropriate to investigate the potential of reverse correlation to probe automatic classifiers, as its advantages and limitations are already well understood for non-linear systems.

One important benefit of the reverse correlation framework is its complete independence from the underlying classifier’s architecture. Unlike efficient but specific methods tuned to a classifier’s architecture (see Guidotti et al., 2018 for a review), the reverse correlation can be used to probe any algorithm that separates the input data into distinct classes. Even for the currently popular agnostic interpretability methods, this is not always the case: Class Activation Mappings (Zhou et al., 2016) are specific to convolutional networks; LIME (Ribeiro et al., 2016) and RISE (Petsiuk et al., 2018) highlight features of specific examples which may or may not be representative of the classification task and stimulus set in general. Also, the method can operate in the same representation space used as an input to the classifier, and can be applied to any type of representation (2D images, 1D time series such as audio, higher-dimensional brain imaging data, for instance).

The outline of the method is as follows. First, a set of stochastic inputs are generated, by introducing noise on the training dataset when available, or, when unavailable, by generating pseudo-random samples designed to cover the input space. The noise takes two forms: multiplicative low-pass noise, known as “bubbles” (Gosselin & Shyns, 2001, 2002), or additive noise, as is more classically the case for reverse correlation. Second, the inputs are sorted according to the classification results. Third, the noise fields associated to inputs correctly classified in the same class are grouped together, with some refinements of the standard reverse correlation methods inspired by signal detection theory (Green & Swets, 1966) to weigh the results with the variability observed after classification. The two variants of input noise, multiplicative or additive, aim to probe two kinds of possibly overlapping but not necessarily identical input features: (1) the *discriminative* features, probed with multiplicative noise, which correspond to the parts of the input dataset that have the most weight in the classification decision and (2) the *represented* features, probed with additive noise, which correspond to the input features most representative of each category. In the machine-learning literature, these would loosely correspond to the “attribution” versus “feature visualization” problems (Olah et al., 2017). In psychophysics, the idea of “potent information” has been introduced in the bubbles framework by Gosselin et al. (2002), for categorization tasks where the ground truth is known. Discriminative features are equivalent to potent features when classification labels for the stimulus class are available, and extend the idea to cases where only the response class is available. The represented information can be compared to the older notion of “prototypical information” for a category (Rosch, 1983).

## 1. Material and Methods

### 1.1. Probing discriminative features

Following the psychophysical literature (Gosselin & Shyns, 2002), we term “discriminative features” the subsets of the input set that are the most positively correlated with the decision taken by the classifier. The aim of this first method is to uncover where such features are located over the whole input dataset. From now on, we assume that the classifier has been trained and is available to the probing method.

#### 1.1.1. Procedure

To identify discriminative features, the input space is pseudo-randomly sampled with multiplicative low-pass filtered noise applied to training samples. A reverse correlation analysis is then applied to identify the noise masks correlated with the classification decision. The algorithm is directly inspired by the “bubbles” method (Gosselin & Shyns, 2001), originally designed to characterize the visual features underlying human behavioral performance for image classification tasks. Having low-pass noise allows to control for the resolution of the probing, enabling a more efficient search with lower resolution. The multiplicative method also focuses the probing to features actually present in the input dataset, further increasing efficiency.

We present two sub-variants of the method, to account for the availability or not of the training set to the end user: 1a) multiplicative lowpass noise is applied to the training set; 1b) multiplicative lowpass noise is applied to noise generated in the input stimulus space. We now describe the algorithm, jointly for 1a) and 1b). The algorithm is illustrated in Figure 1. A software repository written in the Python programming language is provided, including code for all of the examples presented, and available at https://github.com/EtienneTho/proise.

**Fig. 1.**
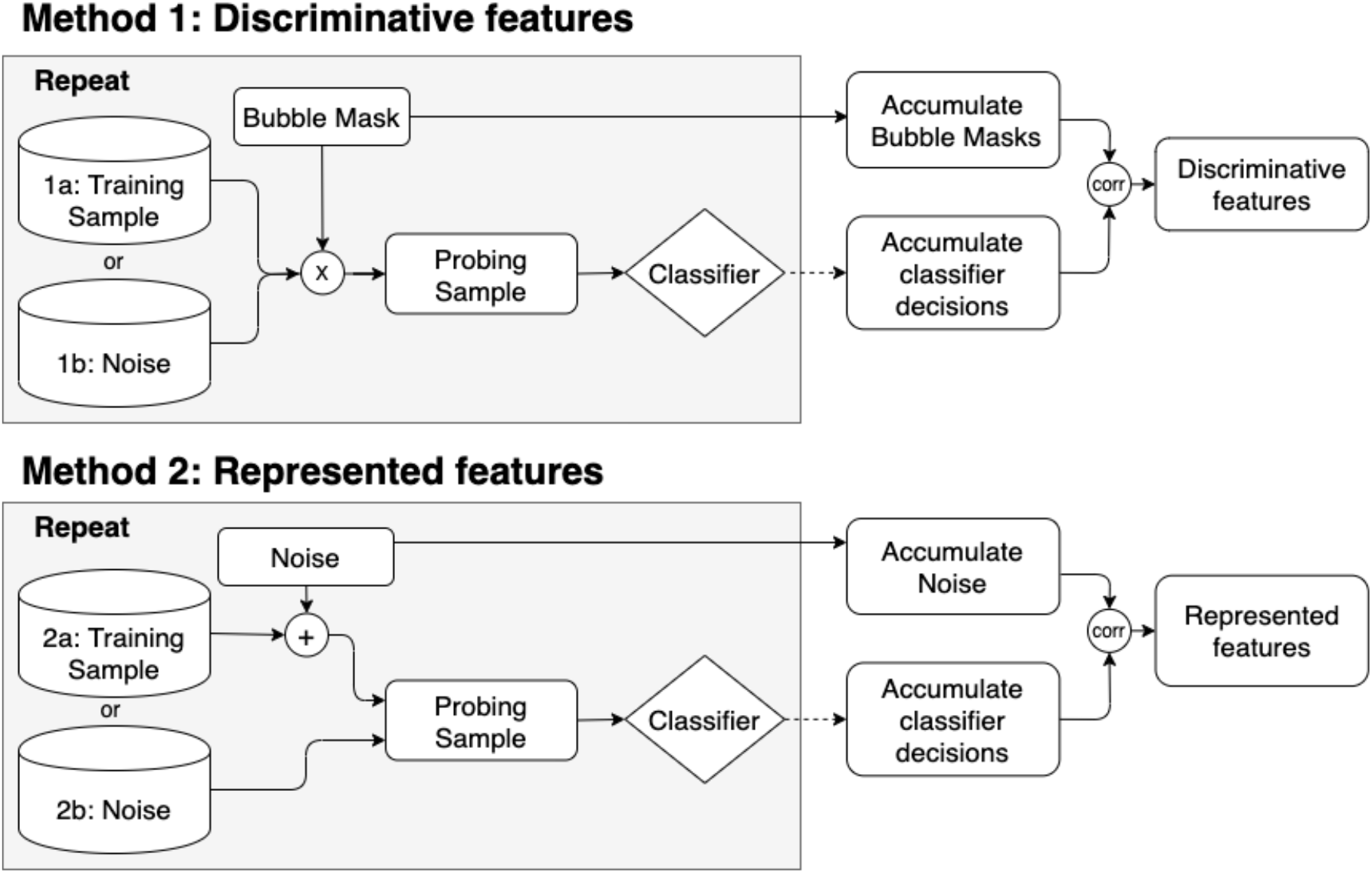
Summary of the two probing methods. Both methods have two variants depending on the availability of the training set. **Method 1: Discriminative features:** Training samples (1a) or noises (1b) are multiplied by bubble masks to obtain the probing samples that are fed to the classifier. The vectors of bubble masks values at each point of the input space are then correlated with the vector of classifier’s decisions to obtain the discriminative features map. **Method 2:** Training samples combined with additive noise (2a) or simply noises (2b) are used as probing samples fed to the classifier. The vector of noise values are then correlated with the vector of classifier’s decisions to obtain the represented features map. For methods 1a) and 2a), only correct classifications are considered, so that the reverse correlation is relative to the stimulus class. For methods 1b) and 2b), there are no labels to the noises and no correct or incorrect classification, so the reverse correlation is relative to the response class. The noise distribution is arbitrary but can be derived from a limited set of unlabeled testing samples (see text for details). Also, in all cases, the classifier’s decisions can be expressed as the binary class decision or as continuous probabilities of class belonging when available.

We now provide a textual description of the main algorithm and its variants.

For each pass:

1. A bubble mask is generated. This consists of a mask in the input space, of dimension N, consisting of randomly positioned N-dimensional Gaussian windows called bubbles (see Figure 2). The number of bubbles is defined probabilistically, to avoid spurious correlations between the presence of a bubble in one part of the representation with the absence of bubbles in other parts of the representation^1^. In practice, the probability that there is a bubble at one point in the input space is set by the probability *P*_bubble_ fixed over the whole input space. This probability and the size of the bubbles in terms of the Gaussian standard deviations are hyperparameters of the algorithm, which can be arbitrarily chosen to fine-tune the output resolution. In practice, an input array of dimension N populated by zeroes except for *nbBubbles* unit values is convolved with N-dimensional Gaussian windows. The resulting mask is denoted *BubbleMask*.
2. The probing data is generated. For variant a), the probing data is one exemplar of the training dataset, randomly chosen. For variant b), the probing data is an N-dimensional activation noise (see section 2.1.2 for details and a suggested technique to obtain an appropriate coverage of the input space). The probing data is denoted *ProbingData*.
3. The probing sample is obtained by multiplying the bubble mask with the probing data: *ProbingSample = BubbleMask * ProbingData*.
4. The probing sample is fed to the classifier and the response class is recorded. A vector *C* is created with a dimension corresponding to the number of possible response classes and populated with 0s. For variant a), the probing sample is labeled 1 for its response class if it is classified correctly according to its stimulus class, 0 otherwise. For variant b), the probing sample is always labeled 1 for its response class as there are no labels for the noise samples and thus no correct or incorrect classification. For each probing sample, we therefore obtain a vector *C* of 1s and 0s describing the classifier’s decisions. Alternately, when available, the continuous probability of the probing sample to belong to the target category can also be recorded in *C*.

**Fig. 2.**
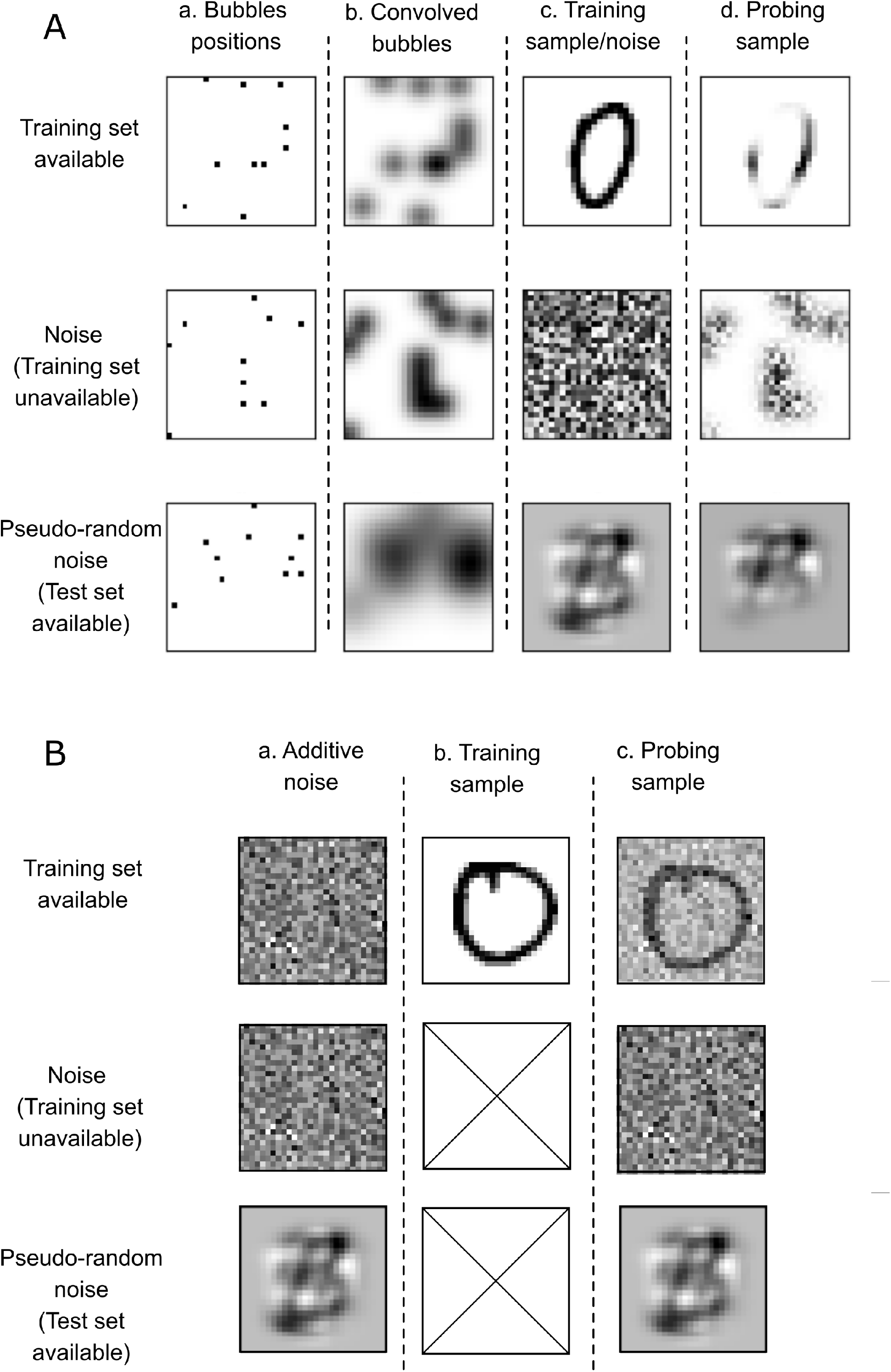
Illustration of the probing method with bubbles and reverse correlation on 2-D images. **Panel A. Top row: construction of the probing samples for discriminative features with the training set available. a)** Random placement of bubbles **b)** Convolution with 2-D Gaussian distributions to obtain the bubble mask **c)** A training sample **d)** Bubble mask applied to the training sample to obtain the probing sample. **Middle row: discriminative features with training set unavailable, uniform noise.** Columns as above, except that the training sample is replaced by uniform noise. The resulting probing samples do not cover the output categories efficiently (see text), likely because of their qualitative differences with digits. **Bottom row: discriminative features with training set unavailable, dimension-reduced noise.** Here the training sample is replaced by noise generated in a reduced dimension space extracted from PCA over the test set. The resulting probing sample is not a recognizable digit but shares visual features with actual digits. Note that it seems that a circular mask was applied to the probing sample, but this is not the case: the lack of features in the periphery is simply a result of the structure of the PCA-based latent space from which the noise was generated. **Panel B. Top row: construction of the probing samples for represented features with the training set available. a)** Generation of the additive noise, here Gaussian noise. **b)** A training sample **c)** Additive noise applied to the training sample to obtain the probing sample. **Middle row: represented features with training set unavailable, noise.** Because training samples are unavailable, the probing sample is identical to the noise sample. **Bottom row: discriminative features with training set unavailable, dimension-reduced noise.** Here the noise sample is generated in a reduced dimension space extracted from PCA over the test set. The probing sample is equal to the noise sample.

Reverse correlation analysis:

For each point *i*, in the input space, the discriminative map for a target, *D*_i,C_, is computed as the correlation between the bubble masks values at point *i, BubbleMask_i_* and the classifier’s decisions *C.* It should be noted that the analysis is performed on the bubble masks, and not on the probing samples themselves. The Pearson or Spearman methods can be used to estimate the correlation coefficient.

#### 1.1.2. Generation of probing samples activations when the training set is unavailable

As mentioned above, if the training set for the classifier is unavailable to the end user, probing samples can be generated from noise in the input space. The choice of the noise distributions is also a free and crucial parameter of the method. The simplest choice is to draw samples from a uniform or Gaussian distribution at each point of the input space, with distribution boundaries covering the full range of valid input values. However, this sometimes leads to uneven coverage of the output categories, for instance if the classifier’s boundaries are especially complex or if the decision algorithm is highly non-linear. Obviously, categories not covered by probing samples cannot be interpreted.

In this case, we suggest to generate pseudo-random probing samples by first randomly inverting the dimensions of the latent space of a Principal Component Analysis (PCA) trained on a representative set of inputs relative to the classifier’s task. These can be obtained for instance from an unlabeled test set. There are few use cases where no exemplars from test set are available, as the test set basically defines the task of the classifier. After the PCA, noise can be generated in the low dimension space and inverted to obtain noise in the input space. The important point here is that, in a whitened representation such as a PCA, it is possible to systematically sample the latent input space by sampling each PCA dimensions with unit-normalized Gaussian noise. Conversely, it is more difficult to homogeneously sample the original feature space without any a priori knowledge of its latent structure.

This method can also be useful when variant a) returns a noisy or uninterpretable discriminative map, even if the training set is available. Indeed, it is only the noise field that is averaged in the reverse correlation approach, and as a consequence some *a priori* choices of noises can be particularly inefficient if they do not match roughly the statistics of the stimulus set, or if the dimensionality of the problem is high. The choice of the noise distribution is thus a free parameter, with Gaussian noise an obvious a priori choice (Gosselin & Shyns, 2002). However, handcrafted optimization of noise has also been proposed for the classification of visual images, for instance (Brinkman et al., 2017). Our PCA-based method, in contrast, is a general and data-driven approach to tailor the noise to the stimulus set statistics, to improve the efficiency of the reverse correlation approach^2^.

In any case, the qualitative goal during the generation of probing samples is to achieve a balanced coverage of all output categories after classification. Iterative choices when generating such probing samples, using a PCA as just described or directly from the raw input space or from any other method, may actually participate in the understanding of the classifier’s features.

#### 1.1.3. Statistical testing

The discriminative features maps illustrate, in the input space, the location of the features used by the classifier to assign samples to a given category. Visual inspection may be sufficient to get a qualitative understanding of the classifier’s operation. However, in some cases, it may be desirable to assess statistically the relevance of each feature appearing in the maps.

There are many options to assess the significance of such data, from which we outline here one possible methodological choice. First, the maps can be shuffled by running the algorithm described above many times while randomly assigning output categories to each sample. For each point in the actual map, a *t*-test (or a non-parametric equivalent) is applied to compare the map value with the mean of the shuffled data. Maps are usually high-dimensional so a correction for multiple comparisons is recommended. Again, several standard choices exist, including Bonferroni correction, cluster permutation (Maris & Oostenveld, 2007), or False Discovery Rate (FDR) (Benjamini & Hochberg, 1995). Specific statistical methods have also been derived in the psychophysical literature using the bubbles and reverse correlation methods, based on random field theory (Chauvin et al., 2005) or maximum statistic method (Holmes et al., 1996). Such methods would be especially suited to the evaluation of discriminative or represented features maps. As the application of these statistical procedures is well documented, in the illustrative examples discussed below we focus on the description of our novel method and only provide the raw feature maps, without statistics.

### 1.2 Probing represented features

We define “represented features” as the parts of the input space that would be the most representative of a given class. We estimate them by using additive noise on the input, followed by reverse correlation, in a way reminiscent of receptive fields measurement in neurophysiology or decision template estimation in psychophysics.

#### 1.2.1 Procedure

To build the represented features map, the input space is randomly perturbed with additive noise. The aim is to probe the whole of the input space, even for features that are not represented in the training dataset, with the highest possible resolution. Then, all probes classified as members of the same category are combined, in a direct adaptation of the classic reverse correlation method. Here, we introduce two differences with classic reverse correlation. This would require to know the training dataset or to have a large labeled test dataset. Again, we propose two sub-variants of the algorithm depending on the availability of the training dataset: a) noise in the input space is added to the training set; b) noise is generated in the input space. As for Method 1, the distribution of the noise is a free parameter that can be chosen as discussed in section 1.2.2. The algorithm is illustrated in the bottom part of Figure 1.

We now provide a textual description of the algorithm and its variants.

For each pass:

1. The probing sample is generated. For a), the probing sample is one randomly chosen exemplar of the training dataset, with noise added. The goal is to perturb the input to introduce variability, so that only the most salient information (to the classifier) remains in the reverse correlation average. For b), the probing sample is an N-dimensional noise. The probing sample is denoted *ProbingSample*.
2. A vector *C* is created with a dimension corresponding to the number of possible response classes and populated with 0s. For variant a), the probing sample is labeled 1 for its response class if it is classified correctly, 0 otherwise. For variant b), the probing sample is always labeled 1 for its response class as there are no correct or incorrect classification. As for discriminative features, continuous probabilities may be used instead of binary class decisions.

Reverse correlation analysis:

For each point, *i*, in the stimulus space, the represented map for a target *D*_i,C_, is computed as the correlation between the additive noise array values at point *i* and the classification decisions vector *C*.

#### 1.2.2 Dimensionality reduction

Depending on the architecture of the classifying pipeline and input space, the estimation of the represented map with reverse correlation can be resource-intensive. A standard technique to improve efficiency when training a classifier is to reduce the number of dimensions of the input space, for instance by using PCA. (e.g., Patil et al., 2012). Here too, the probing and reverse correlation analysis can be performed in a space with reduced dimensionality before inverting back to the original input space.

For the generation of probing noise in variant b), the same method described in section 2.1.2 apply, with the potential use of PCA to shape the noise for a balanced coverage of all output classes.

#### 1.2.3 Statistics

The statistical analysis of represented maps can be done with the same generic tools as for discriminative features maps, described in section 2.1.3.

## 2 Results

To illustrate the methods introduced above, we present three different use cases covering different types of input data (2D images, 1D speech and music sounds, multi-channel electrophysiological data) and decision type (binary versus multiclass decisions). First, we interpret the classification of handwritten digits, using a visual task (2-D pixel input space) performed with a convolutional deep neural network. Second, we interpret the classification of speech versus music, using an audio task (1-D audio time series converted to a 4-D auditory model) performed by a support vector machine. Third, we interpret a sleep scoring algorithm aiming at classifying electrophysiological data into sleep stages (3 classes) using 75 spectral features extracted from 30-s time epochs fed to a random decision forest classifier.

Although voluntarily simple, the first two examples are used to introduce the main features of our method. The last example illustrates an application of our method to a concrete neuroscientific problem, as sleep stages classification from polysomnographic recordings is a time-consuming task that would greatly benefit from reliable and interpretable automation (Andrillon et al., 2020). A benefit of our interpretation method is its generality, so the examples covered here are also intended to showcase the various ingredients needed for many use cases relevant to neuroscience.

### 2.1 Digits classification

#### 2.1.1 Classification task

In this first example, we classify visual samples of handwritten digits from the *MNIST* database (Deng, 2012). This is a standard database for evaluating image classification algorithms in the machine-learning community. It is composed of handwritten digits, from 0 to 9, with 60000 samples in the training set and 10000 samples in the test set. Each sample is a two-dimensional greyscale image with pixels values between 0 and 1.

Many algorithms can now successfully perform this classification task. Here we trained a Convolutional Neural Network (CNN) to discriminate between digits, with the following architecture: 2D-convolutional layer (3, 3), Max Pooling layer (2, 2), 2D-convolution layer (3, 3), flattening layer, dense layer with 10 outputs and a softmax activation. Three training iterations were run. As expected, a high classification accuracy of 97% was obtained on the test set.

#### 2.1.2 Discriminative features maps

We first describe the probing of the trained CNN using Method 1, aimed at uncovering the discriminative features maps. We compared the cases for which the training set was available or not, and used the PCA-based method introduced in section 1.1.2 to generate probing samples when only the test set was available. Figure 2 illustrates the kind of probing samples obtained in these various cases.

To estimate the maps, 200,000 probing samples were generated for Method 1a, which is an empirical trade-off between computation time and stability of the results. For Method 1b), we observed that less probing samples were necessary, so ten times less (20,000) were generated. In all cases, a 30% probability to have a bubble at each location was applied. The standard deviation of each bubble was 2 pixels. These choices were also empirically made to achieve good coverage of most classes and an accuracy. With these parameters, the accuracy of the classification during probing was 65.83%. For Method 1b, we used the continuous probability belonging to a class obtained at the output of the CNN to correlate with the bubble masks. This implementation should be favored when the continuous probabilities are readily available. The probabilities were obtained with the function ‘predit_proba’ from the Tensorflow/Keras distribution (Chollet et al., 2015).

Figure 3 shows the resulting discriminative features maps, expressed as the correlation between the bubble mask and the classification decision in the 2-D input space (the visual image). The maps are intended to highlight regions of the input representation that are weighed most in the classification decisions. These regions do not necessarily correspond to the actual shape of a digit (which will be targeted by represented features later on), for two reasons. First, the bubbles were chosen here with a relatively low resolution. Second, bubble masks were multiplied with actual input samples before being fed to the classifier, but the features of these samples are not taken into account in the reverse correlation as it was performed on the masks themselves.

**Fig. 3.**
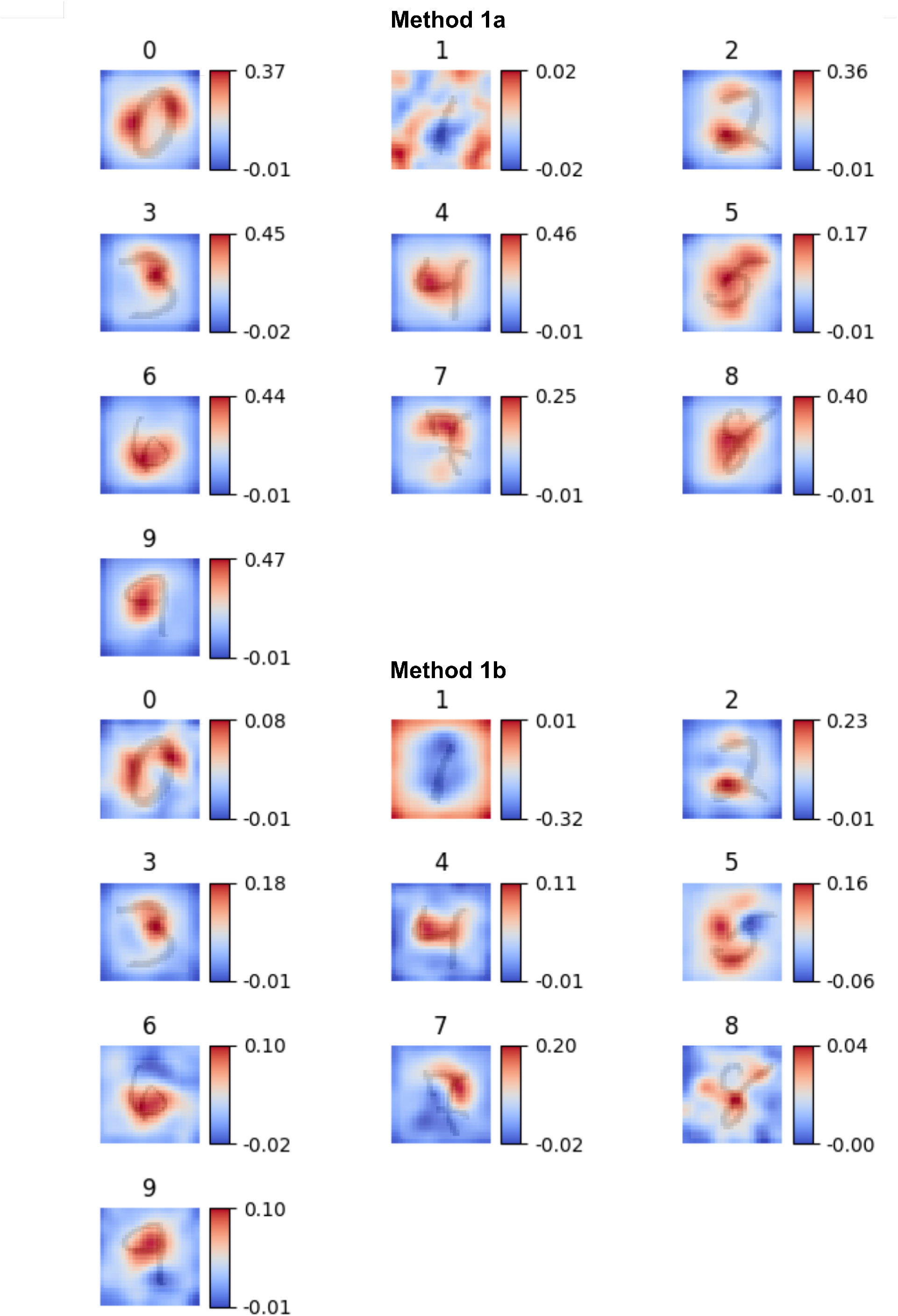
Discriminative features maps for a CNN classifying handwritten digits. Each panel shows the discriminative features for a given response class (digit), expressed as correlation coefficients in the original 2-D input space. To facilitate interpretation, a typical test sample has been overlayed (dark shade) on each map. The top panels correspond to Method 1a with the training set available and the bottom panels correspond to Method 1b with the training set unavailable. Regions in red highlight the locations of the input space for which active pixels are associated to classification in the digit’s class.

For some digits, such as “2” and “7”, the maps highlight part of the stimulus that are likely unique to each digit: a loop in the bottom left corner for “2”, sharp corners in the top part of the image for “7”. These features are most clearly visible in the output of Method 1b. Interestingly, for the digit “1”, most of the positive correlations appear at the periphery of the map whereas negative correlations are where the digit image should be: knowing that there are no active pixels in peripheral regions appears to be the most efficient cue for deciding that the uniquely narrow shape of “1” was the input.

In summary, while the discriminative features maps may not make immediate intuitive sense on their own, they do orient the analysis of the input set towards regions of interest, especially when interpreted along representative input samples as illustrated in Figure 3. Moreover, if the task was now to classify e.g. “2” versus all other digits, then the input space could be weighed to emphasize the bottom-left corner to simplify the new classifier.

We now compare more formally the method’s outputs when the training set is available or not. The availability of training data is expected to provide faster and more robust convergence towards the features of interest. However, this was not the case here, as we needed to use less probing sample for Method 1b compared to 1a to obtain less noisy discriminative maps. This is because we used pseudo-random noise in a PCA-reduced space obtained from the test set as probing noise for Method 1b (see section 1.1.2). An initial attempt with simple Gaussian noise led to unbalanced classification and no decision in four digit categories, which prevented interpretation of these categories. We then used 10^4^ test set samples to build the PCA space, retaining 100 dimensions for the PCA. Finally, 2.10^4^ probing samples were then generated using normalized Gaussian noise in the PCA space and inverting back to the 2D original input space. This stochastic sampling procedure led to decisions covering the 10 categories in a more balanced fashion. A systematic investigation of the effect of these three parameters (number of test samples, PCA dimensions, number of probing samples) is provided in Supplemental Figures 1 and 2.

In any case, as can be seen from Figure 3, the discriminative information obtained with Method 1b was highly similar to that uncovered by Method 1a. This was confirmed by the strong correlations between maps across these methods with or without training set available (average Pearson coefficients for the 10 maps: M = .80 (SD = .16), *df* = 783, all *p* < 10^−3^).

#### 2.1.3 Represented features maps

We now probed the same trained CNN with Method 2, aimed at uncovering represented features maps. Such maps are can be interpreted as weighted averages of probing samples themselves, obtained by reverse correlation between the CNN’s decisions and the input features. We used 250000 probing samples for Method 2a and 10000 probing samples for Method 2b, again following an empirical trade-off. For Method 2a, we used additive Gaussian noise as a naïve a priori choice, but other type of noises can also be used (Brinkman et al., 2017, see also Supplemental Figures 3 & 4).

Figure 4 shows the resulting represented features maps. Qualitatively, these maps look very different from the discriminative feature maps. Figure 2a in particular looks especially noisy, with the additional issue that the class for the digit “1” was never observed and could not be sampled. This is because the probing samples were directly generated in the input space, without any low-pass filtering, allowing the expression of finer details but also impeding convergence. Also, for a correct classification of the digit “1”, as was observed in the discriminative map, there should be no energy in peripheral regions of the image. Such a case never occurred with unconstrained Gaussian noise sampling, which is why the method failed to generate a map for this class. Gaussian noise is not the only possible choice of noise, however, and studies of image classification tasks have suggested the use of Gabor (Brinkman et al., 2017) or Fractal noises (Hansen et al., 2010). Such noises indeed reduced the noise in the represented features maps (see Supplemental Figures 3 & 4).

**Fig. 4.**
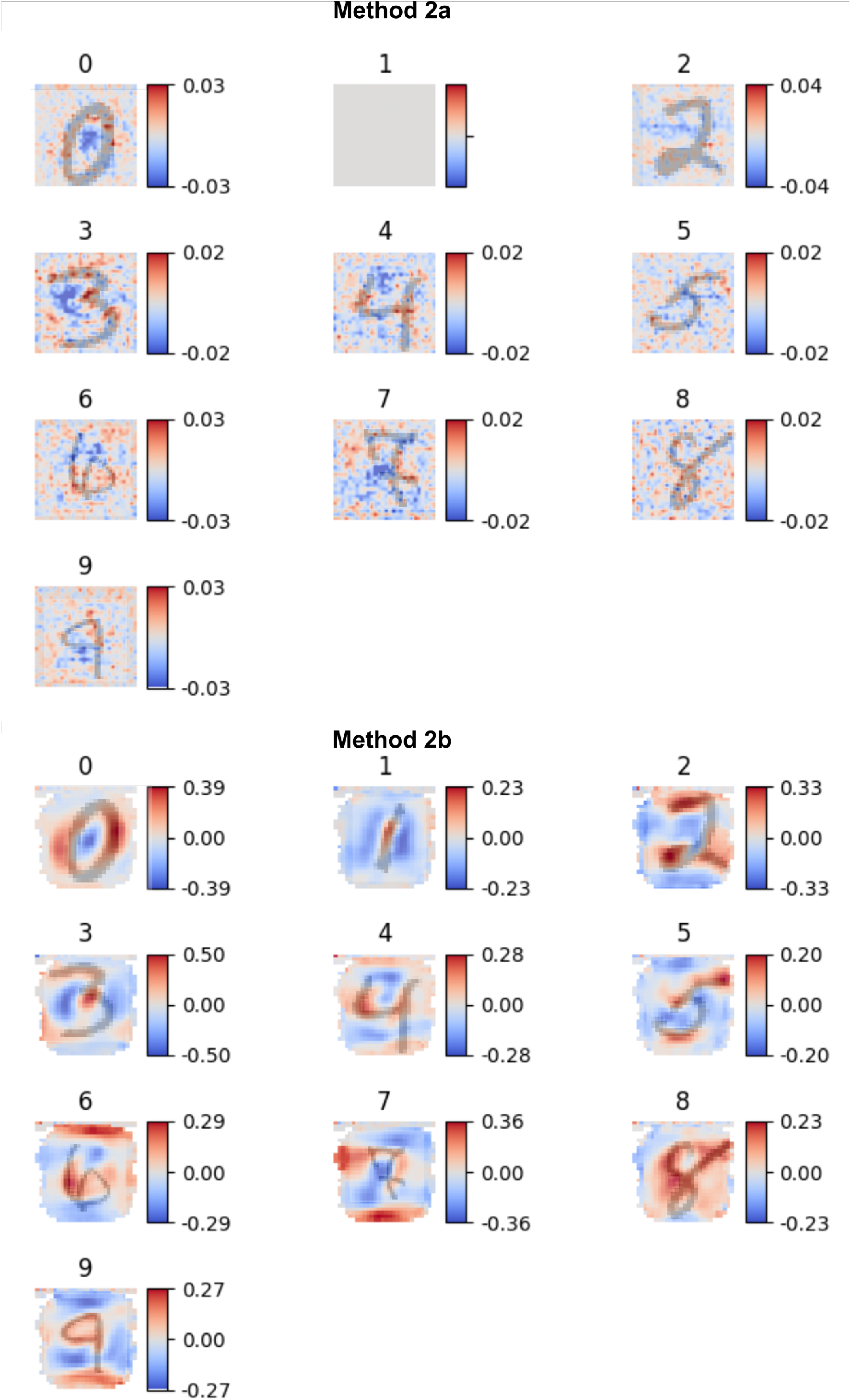
Represented features maps for a CNN classifying handwritten digits. The different panels show the represented features for each stimulus class (digit), expressed as correlation coefficients in the original 2-D input space.. To facilitate interpretation, a typical test sample has been overlayed (dark shade) on each map. The top panels correspond to Method 2a with the training set available and Gaussian noise while and the bottom panels correspond to Method 2b with the training set unavailable and PCA-based noise. For Method 2a, no probing sample was ever correctly assigned to the digit “1” so no map could be computed. The red portions of the maps indicate the input features most associated with a given class. For Method 2b at least, they qualitatively resemble each digit. The blue portions of the maps indicate the input features that are most reliably not present for a given class

The maps look less noisy and more interpretable for Method 1b), with the PCA-based noise used as probing samples. As expected with a reverse correlation approach, in this case the represented features maps match the “best” stimulus of a category. The category of each map is readily identifiable, as it visually resembles the written digits’ representations. In addition, ‘negative’ regions in blue further specify which features are expected to be absent to classify a given digit. Altogether, here the represented features maps behave as expected in a digit classification task, and without any information about the classifier, it was able to recover representative digit-like shapes for each category.

In spite of the different amount of noise visible in the maps, Methods 2a and 2b actually provided correlated maps (average Pearson coefficients for the 10 maps: M = .29 (SD = .11), *df* = 783, all *p* < 10^−3^). This confirms that Method 2a uncovered essentially a noisier version of the maps. It can nevertheless be noted that Method 2b tended to focus on the center of the input space. In particular, some border pixels were never associated with one or the other classification decision, leading to missing values when computing the correlations. That is because the noise itself was shaped by the stimulus-derived statistics, which led to less probing on the periphery.

#### 2.1.4 Quantitative evaluation

So far, we have qualitatively described the output of the probing methods, and used such descriptions to gain some insights on the classifier’s decisions, as would be the case in realistic use cases. However, as mentioned in the Introduction, there are already established interpretation methods for some classifiers’ architectures. In particular, as we used a CNN for the digit classification task, we could compare the output of our algorithms to the Class Activation Mapping method (CAM, Zhou et al., 2016), a state-of-the art technique to visualize the features of a trained CNN.

To do so, we first adapted a metric proposed by Petsiuk et al. (BMVC, 2018) to evaluate the relevance of the visualized information. The gist of the metric is to evaluate the importance of features present in an interpretation mask by systematically adding or removing features from the mask. If the mask captured relevant features, then adding mask features should improve classification, whereas removing mask features should impair classification. Here, we used the discriminative features maps as interpretation masks.

In practice, the masks were first binarized by *z*-scoring the distribution of each mask and attributing the value 1 to the pixels which were above the 0 and 0 otherwise. Then, for each sample of the test set, we added or deleted a proportion of features in the mask, i.e. mask values were randomly flipped from 1 to 0 or the reverse. Masks were then blurred again, as in Petsiuk et al. (2018). This was achieved through a convolution with a 2D Gaussian window (standard deviation of .3 pixels). Finally, masks were multiplied with testing samples. For each given proportion of insertions or deletion, the accuracy of the classifier was tested and the Area Under the Curve (AUC) of the accuracy vs. proportion curve was used as the performance metric. In the case of the insertion, the higher the AUC, the better the interpretation. Conversely, in the case of the deletion, the lower the AUC, the better the interpretation.

We applied this method on all of the samples of the test set for the MNIST task. For insertion on discriminative features maps, we observed a mean AUC=.75 (SD=.23). For deletion on discriminative features maps, we observed a mean AUC=.14 (SD=.31). The difference between the two values was highly significant (two-sided t-test, t(19998)=158.58, p < 10^−5^, d_cohen_=2.24), demonstrating that the discriminative features were relevant for the classification task.

We then generated interpretation masks using the CAM method (Zhou et al., 2016). We used the default parameters of a grad-CAM implementation (https://raghakot.github.io/keras-vis/vis.visualization/#visualize_cam), selecting the gradient based class activation map (grad-CAM) that maximizes the outputs of all filters in the last convolution layer of the CNN as the mask being tested for each sample of the test set. From this mask we then applied the same steps as for the bubbles method to compute the insertion and deletion AUC metrics, i.e. thresholding and blurring. For insertion with a mask obtained with CAM, we observed a mean AUC=.56 (SD=.29). For deletion, we observed a mean AUC=.19 (SD=.30). The difference between the two values was highly significant (two-sided t-test, t(19998)=88.29, p < 10^−5^, d_cohen_=1.25), showing that the CAM masks also captured relevant information for the classification task, as expected. However, the difference between insertion and deletion AUC was smaller for the CAM masks than for the discriminative features masks (two-sided t-test, insertion: *t*(19998)=-51.03, p < 10^−5^, d_cohen_=.72; deletion: t(19998)=10.91, p < 10^−5^, d_cohen_=.72). This suggests that our method outperformed CAM for the quantitative metric chosen and the example tested here.

### 2.2 Speech vs. music

In this second example, we classified audio samples in a speech versus music task, to vary the input space and classifier’s architecture. We used the GTZAN database composed of 132 excerpts of speech and music (Tzanetakis & Cook, 2002). The database was preprocessed to create samples with a fixed duration of 5 seconds, leading to a dataset of 768 samples. Those samples were randomly separated into a training set (691 excerpts) and a test set (77 excerpts, 10% of the dataset).

Following Patil et al. (2012), who performed an automatic classification of the musical timbre of short audio samples, sounds were first processed by an auditory model (Chi et al., 2005). The idea was to cast the input space into a representation that is interpretable in terms of auditory processing, unlike the raw waveform representation. Briefly, a filterbank corresponding to cochlear tonotopy was initially applied, followed by a 2-D Fourier analysis of the resulting time-frequency representation. The model output thus represents temporal modulations and spectral modulations contained in the input sound (Chi et al., 2005, Elliot & Theunissen, 2009). The 4-D resulting arrays, with dimensions of time, frequency, scale of spectral modulations, and rate of temporal modulation, are termed here Spectro-Temporal Modulation representations (STM). We averaged the time dimension over the 5s of each sample. Next, we applied a PCA to reduce dimensionality (30976 dimensions in our implementation: 128 frequency channels x 11 scales x 22 rates, reduced to 150 dimensions to preserve 98% of the variance).

For classification, the output of the reduced PCA was fed to a Support Vector Machine (SVM) with a Radial Basis Function (RBF). All of these steps are identical to Patil et al. (2012), to which the reader is referred to for further details, as the specifics of the classifier are not critical to illustrate the probing method. Briefly, a grid search on the RBF was performed to determine the best set of parameters and the classifier accuracy was tested with a 10-fold cross-validation. We obtained an average classification accuracy, i.e. whether the classifier is classifying the STM of a sound to the correct music or speech class, of 94% (SD = 6%) with the 10-fold cross-validation and 98% on the test set.

Figure 5 shows the discriminative feature maps for the speech versus music classification task. For each case, we used 10000 probing samples and a probability of 0.3 for the bubbles with standard deviation of 10 Hz in the frequency dimension, 5 Hz in the rate dimension, and 5 cycles/octave in the scale dimension. With these empirically-determined parameters, the accuracy of the classification during probing was 80.8%. As the task was a binary classification, the maps for speech and music were simply mirror images of each other. We only present the maps corresponding to speech (the reddest parts) vs. music (the bluest parts). Also, in order to test another implementation of the analysis framework, correlations were obtained using the binary decisions of the classifier and not continuous probabilities.

**Fig. 5.**
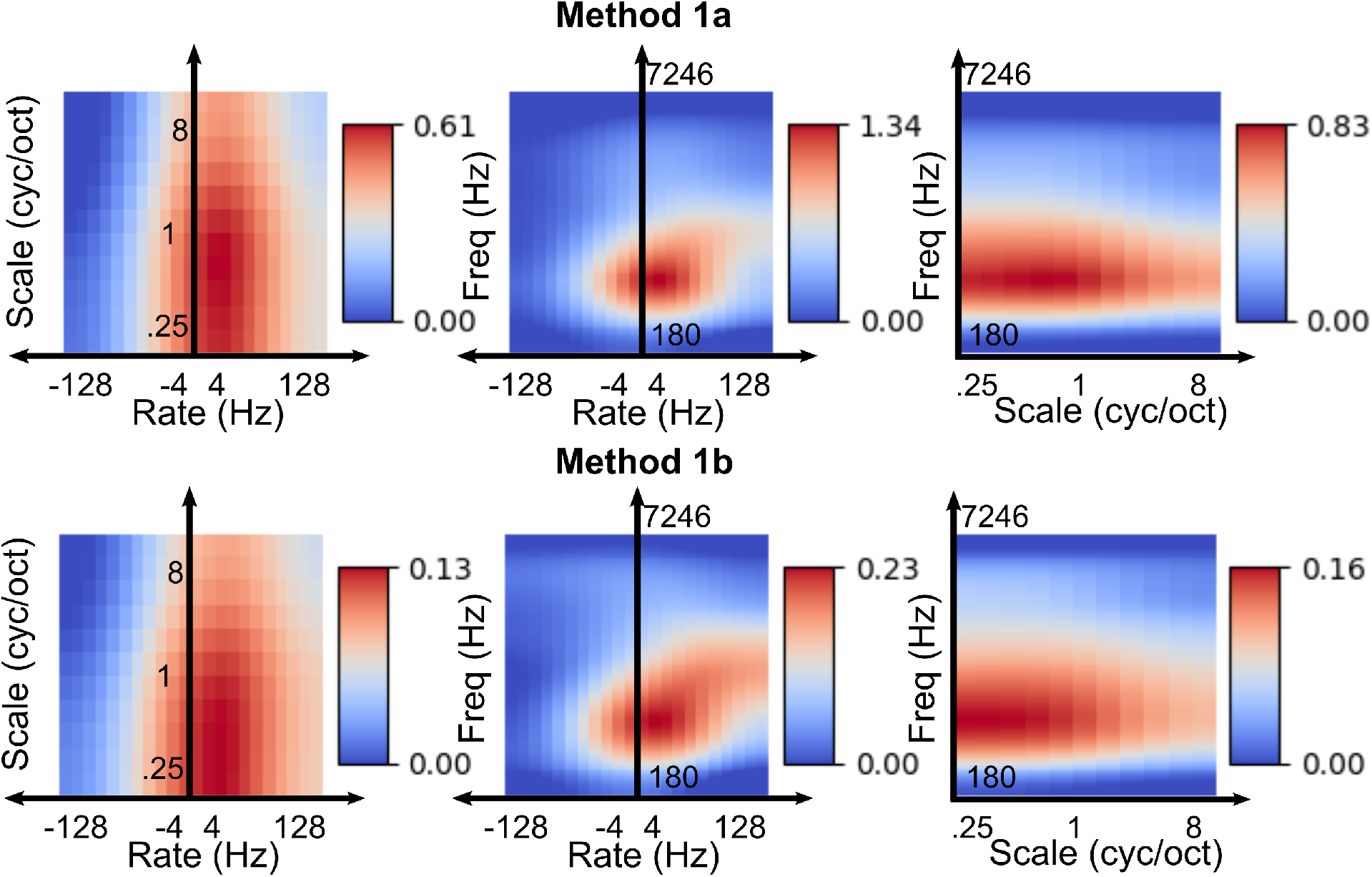
Discriminative maps for speech and music in the STM representation. The 4-D auditory model representations are averaged over time and projected in the remaining three dimensions (frequency, scale, rates). Maps are expressed as correlation coefficients between the bubble masks and the binary classifier’s decision. Method 1a uses the training set while Method 1b uses pseudo random noise. As this is a binary classification task, the speech and music masks are simply opposite versions of each other.

The discriminative regions of the auditory model STM representation appear to be mostly visible in the frequency dimension: speech can be best classified by looking at the input in a broad frequency range around 500 Hz, corresponding roughly to the position of the first formant in speech (Peterson & Barney, 1952). For the other dimensions, the classification depends on slow positive rates and low scales. In other words, the difference between speech and music was in the presence of slow modulations and broad spectral shapes for speech. Again, this matches prosodic and syllabic features of speech, together with the broad spectral shape of formants. By construction the two maps are complementary, but for music, a richness in spectrum, including high and low frequency regions, associated with fine spectral details (high scales) is characteristic of musical instruments (Elhilali, 2019), which have been designed to go beyond the physical constraints imposed by voice production. This is consistent with the blue parts of Figure 5.

In the case of the Method 1b, with the training set unavailable, we used pseudo-random noise generated in a PCA-reduced representation obtained with the test set. A PCA was first applied to the test set to reduce it to 50 dimensions and a uniform Gaussian white noise was generated on the 50 dimensions to generate samples in the reduced space. Each random reduced sample was then transformed into the original input space by applying the inverse PCA transformation. This procedure allowed to generate noisy samples with distribution relevant regarding the representative set of data relevant to the classification task. The information obtained by the two methods strongly correlate (Pearson coefficient: *r* = .95, *df* = 30974, *p* < 10^−5^), even though differences are visible on the frequency dimension.

Figure 6 shows the represented feature maps for the speech versus music classification task. The represented features maps provided by Method 2a were extremely noisy and uninterpretable, even more so than for the digits example. We hypothesize that this has to do with the naïve choice of noise and the high dimensionality of the problem (36976 features in the STM representation). However, Method 2b again provided interpretable maps, even though it used less probing samples than Method 2a (10000 against 30000). Compared to the discriminative features maps of Figure 5, these represented features maps show in finer details what a speech sound or a music sound may look like in the STM representation. Similarly, they indicate the “average” speech and music sounds learnt by the SVM. For speech, some formantic structure is now visible on the frequency dimension, associated with low rates typical of prosodic modulations (middle panels). These formantic regions extend to higher scales (right panels), perhaps because formants are superimposed on a harmonic structure for vowel sounds. Conversely, musical sounds more typically contain high modulation rates and spectral scales. These observations are consistent with previous analyses of STM representations (Elliott & Theunissen, 2009; Chi et al., 2005).

**Fig. 6.**
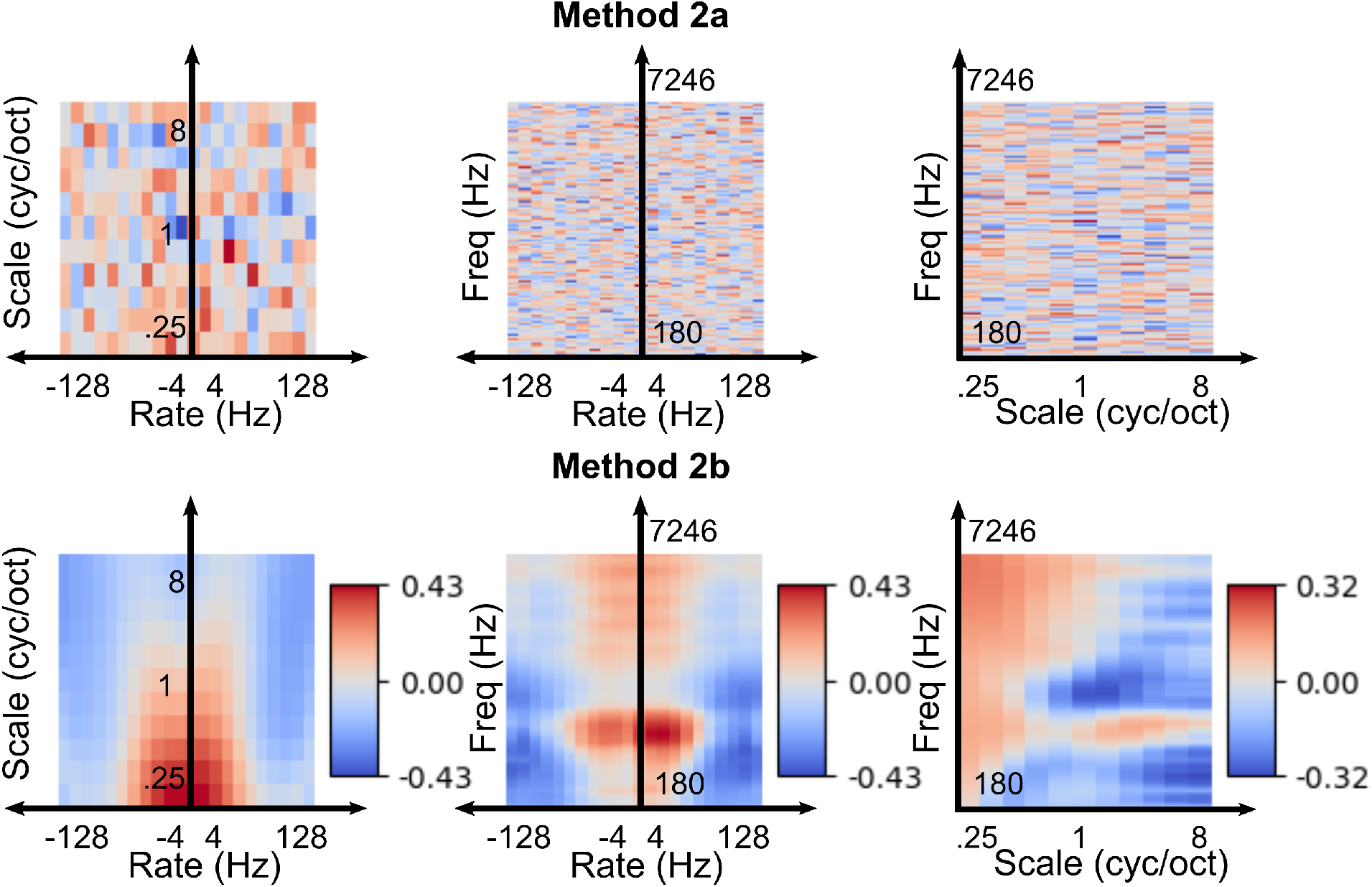
Represented STM representations for speech and music. The 4-D STM representations are averaged over time and projected in the remaining three dimensions (frequency, scale, rates). Details as in Figure 5. The method 2a failed to provide a relevant information in this case.

It should be noted that represented features depend on the acoustic characteristics of speech and music, but they also depend on the task of the classifier. Probing a classifier trained to discriminate speech from e.g., environmental sounds would likely provide different represented features for speech. This result may seem like a limitation of the method, but it also highlights the way an automatic classifier performs a binary task. This may be an important difference to keep in mind when comparing classifiers with human perception, which has to perform many concurrent tasks in parallel. For human perception, “opportunistic features” that depend both on sensory information and the task at hand have been suggested for auditory timbre recognition, a task not unlike the one probed here (Agus et al., 2019).

### 2.3 Classification of sleep stages

In this third example, we interpreted the classification of sleep stages from neurophysiological recordings by a random forest classifier. This represents yet another type of input data and classifier architecture, with a direct link to a real and non-trivial classification problem.

We used the Sleep Physionet database, composed of 153 polysomnographic recordings (PSG) from 78 subjects (41 females and 37 males) (Kemp et al., 2000). From this database, for simplicity, 15 subjects were randomly chosen for the training and 15 others for the testing. Each recording includes 2 electroencephalographic (EEG), 1 electrooculographic (EOG) and 1 electromyographic (EMG) derivations. The data is segmented in epochs of 30s, and each of these epochs have been scored by human experts in 6 categories, according to established guidelines (Rechtschaffen & Kales, 1968): wake, NREM stage 1-4 and REM. There was also 1 additional non-sleep category (movements) and 1 unscored category (total number of original categories: 8). Four of the 6 sleep stages being subdivision of NREM sleep (NREM stage 1-4), we re-labelled the epochs according to the three distinct physiological states: wake, NREM and REM sleep. Movement and unscored epochs were discarded. The database was then preprocessed to extract the Welsh Power Spectrum Density (PSD) between 0.5Hz and 30Hz on 30s samples of the EEG channels (Chambon et al., 2018). We obtained a total of 12932 samples for the training and 13110 for the testing. Each sample had 75 frequency features. We then followed a published classification pipeline (Chambon et al., 2018) by applying first a PCA to reduce the number of features to 30 and then trained a Random forest with a grid search for the best hyperparameters to predict the sleep stage. The training led to a classification accuracy of 99.1% on the training set and 74.9% on the test set, which is not optimal but higher than the chance level of 33%.

Figure 7 shows the discriminative features maps expressed as correlation visualized in the 1-D input space, the EEG power spectrum. For method 1a, 100000 probing samples were used, for method 1b, 10000 probing samples were used. In each case, a probability of .3 to have a Gaussian bubble in each bin, with a standard deviation of 3 bins was used. With these parameters, the accuracy of the classification during probing was 38.72%. Interestingly, the discriminative features reflect what is known of the physiological differences between wake, NREM sleep and REM sleep. In particular, wake led to negative correlation coefficients in the lower frequency range (<5Hz) and positive correlation in the higher frequency range (>20Hz). This is in line with the predominance of fast, small-amplitude desynchronized rhythm and the absence of slow, large-amplitude synchronized oscillations in wakefulness. Conversely, NREM sleep showed positive correlation coefficients for the delta (1-6Hz) and sigma (∼12Hz) range, which correspond to the frequency bands of the hallmark events of NREM sleep: slow waves (1-4Hz) and sleep spindles (11-16Hz). There are not many discriminative features for the REM category, which reflects the fact that REM is best categorized using a combination of EEG, EMG and EOG signals and not just the EEG data considered in this illustration.

**Fig. 7.**
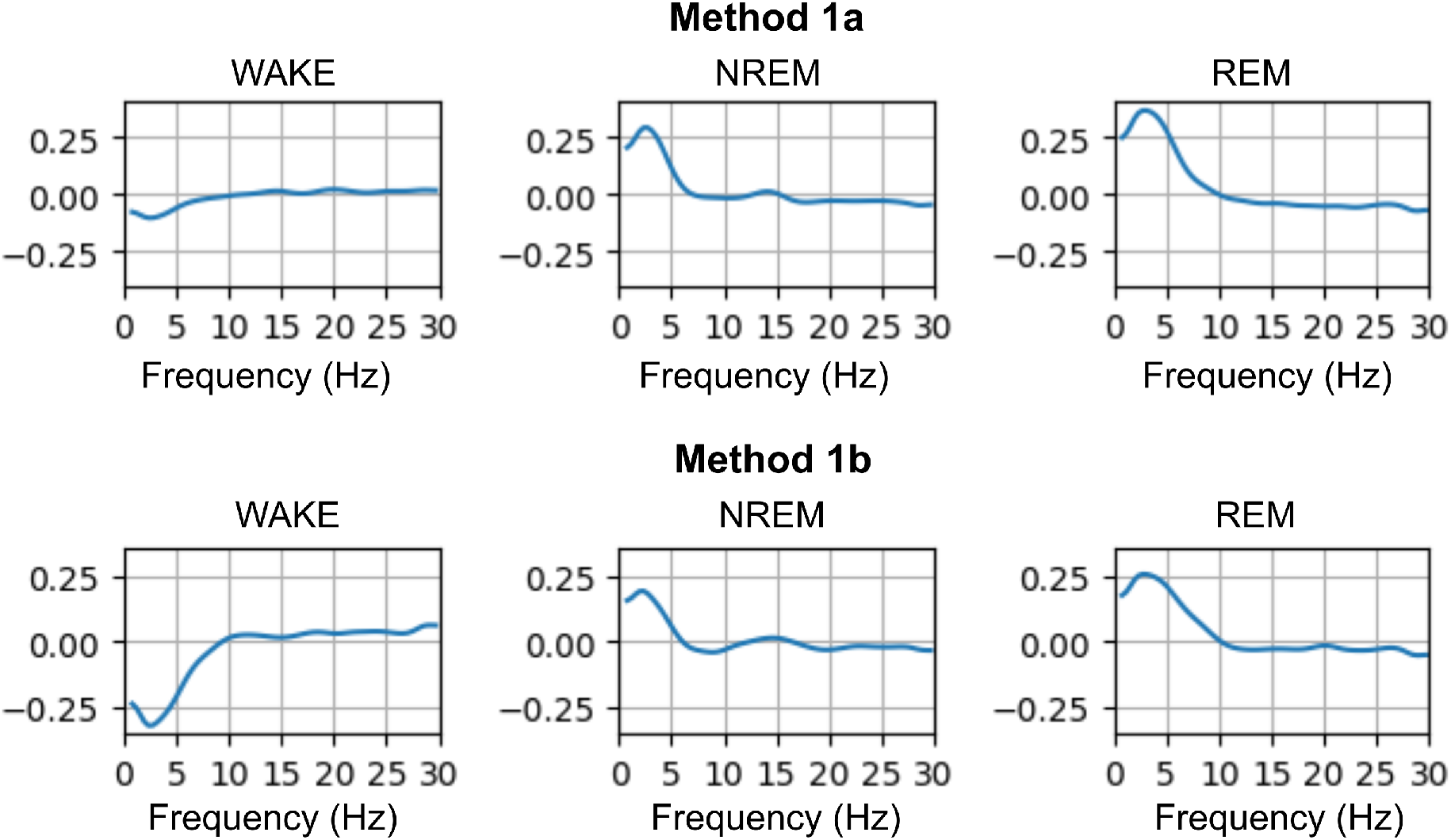
Discriminative features maps for sleep stages classification from polysomnography and a random forest classifier. The panels show the discriminative information expressed as correlations in the input space to the classifier, that is, the power spectrum of the EEG channels of the polysomnographic recording. Positive values correspond to frequency regions where high power is associated to the target sleep stage, whereas negative values correspond to frequency regions where low power is associated to the target sleep stage.

In the case of Method 1b, a pseudo random noise obtained from the inversion of a 30-dimension PCA trained on the test set was used to probe the classifier. The discriminative information obtained in the two cases correlate strongly (average Pearson coefficients for the 3 maps: M = .98 (SD = .01), *df* = 73, all *p <* 10^−3^).

Figure 8 shows the represented features obtained for the different sleep stages. 1311000 probing samples were used were used for Method 2a and 10000 for Method 2b. Interestingly here, Method 2a seemed to converge to interpretable maps, even if it required more probing samples than Method 2b. We interpret this difference with the two previous examples, digits and speech vs music, with the lower dimensionality of the problem here. In terms of the interpretation of the maps, represented features appear even more aligned with the physiological signatures of each sleep stage. Indeed, quiet eyes-closed wakefulness is characterized by predominant alpha oscillations (8-11Hz) and the represented features for wake shows a peak within this frequency band. NREM is characterized by the presence of slow waves (1-4Hz) and sleep spindles (11-16Hz) and the represented features show again positive coefficients for these frequency bands. Finally, in addition to the absence of wake and NREM features, REM sleep shows a positive correlation in the theta band (4-7 Hz), especially for Method 2b, which is in line with the modest increase in theta power observed in REM sleep. However, since REM sleep is not only characterized at the level of the EEG but also with EOG and EMG signals, which were not included in the classifier, the represented (and discriminative) features for REM sleep unsurprisingly show less frequency-specific effects compared to wake or NREM sleep. It is interesting to observe that the represented features map provides somewhat different information than the discriminative features map. This reinforces the idea that the two methods can be complementary.

**Fig. 8.**
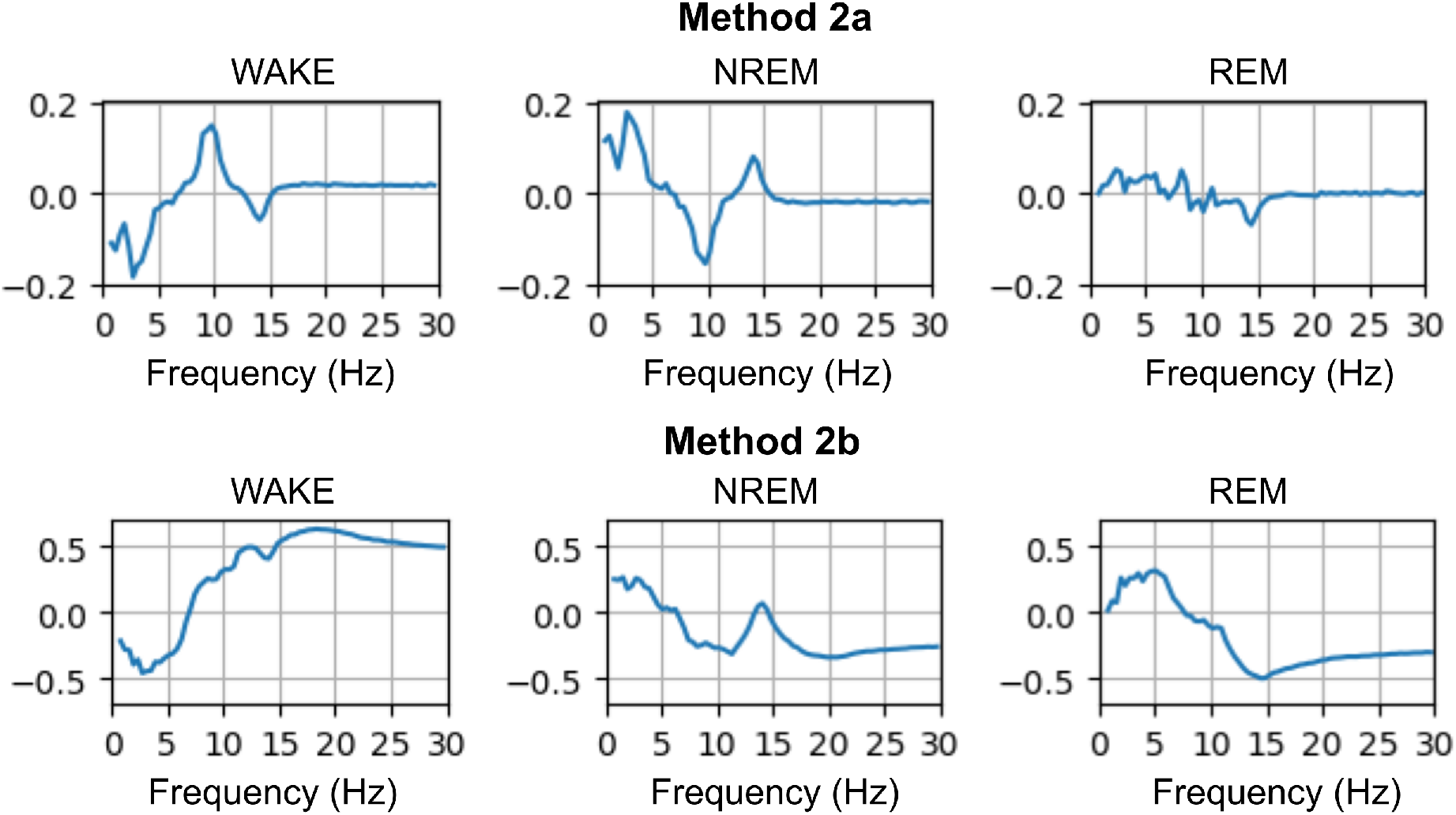
Represented features maps for a random forest classifying sleep stages from polysomnography power spectrum. The plots show the represented information for each point of the input space, obtained with Method 2a (top) and Method 2b (bottom).

The represented information obtained for Method 2a and 2b correlate well stage (average Pearson coefficients for the 3 maps: M = .71 (SD = .08), *df* = 73, all *p <* 10^−3^) although we can note specific discrepancies between the two methods in particular for the wake class.

## 3 Discussion

### 3.1 Summary

The method presented here used a reverse correlation framework to visualize the input features discovered by an automatic classifier to reach its decisions. Over three examples, using different kinds of classifiers (a CNN; an SVM with RBF; a random decision forest) and different kinds of input representations (2-D visual images; 1-D audio samples converted to a 4-D auditory model; multi-channel 1-D EEG data converted to 1-D spectral features), we illustrated how the method could highlight relevant aspects of a classifier’s operation. Moreover, by combining standard noise perturbation techniques with so-called bubbles (Gosselin & Shyns, 2002), we showed that the probing method can be focused either on discriminative features, related to the decision strategy of the classifier, or on represented features, related to the output classes’ main characteristics.

### 3.2 Benefits

In the context of neuroscience and experimental psychology, we believe that there are many benefits in using a reverse correlation framework to interpret classifiers, as a way to complement other more specialized machine-learning interpretation techniques that will undoubtedly remain the preferred tools for purely data-science applications (Zhou et al., 2016; Ribeiro et al., 2016; Petsiuk et al., 2018; Borji & Lin, 2019; Xu et al., 2018).

First and foremost, reverse correlation is a familiar tool in the field of neuroscience and experimental psychology. It has proved useful to gain insights about stimulus features relevant to neural activity, at the single neuron level (Eggermont et al., 1983; Neri & Levi, 2006), network level (Arnal et al., 2015; Adolphs et al., 2005; Ringach & Shapley, 2010), or perceptual decision level (Gosselin & Shyns, 2001; Venezia et al., 2016). Applying it to interpret classifiers amounts to translating and applying a familiar toolbox to another, conceptually similar problem of characterizing a black-box system, this time a computer-based algorithm instead of a brain or a person.

Second, the method is fully agnostic by design. It operates by default on the input space of the classifier, whatever this space might be. It does not make assumptions on the classifier’s architecture or inner operations. Focusing on the input space rather than the classifier’s architecture is especially desirable in situations where the classifier is not the main interest of study, but rather, the structure of the input dataset is.

Third, it can be applied to classifiers that have not been designed by the user, as it does not even require the availability of the training dataset. Access to labeled input data is helpful in improving the efficiency of the method, for instance by allowing to shape the perturbation noise, but this is a mild constraint: there are no interesting situations we can think of for which both the classifier and the type of data to classify would be unknown.

Finally, the output of the method is a visualization (with statistical evaluation if required) in the input space. Such a representation should make intuitive sense to the user, and the features discovered can be interpreted *a posteriori* in terms of attributes of the stimuli. If the representation does not make intuitive sense, then one possible benefit of the method is to help re-cast the input space into a more meaningful representation, as was done here in the audio example for which the waveform samples were pre-processed with an auditory model. This idea is further detailed in the “Perspectives” subsection.

### 3.3 Limitations

There are also limitations associated to the use of a reverse correlation approach to interpret automatic classifiers. Broadly speaking, these limitations follow those already described for reverse correlation in neuroscience.

First, reverse correlation is inspired from the analysis of linear systems, whereas machine-learning classifiers often rely on a cascade of non-linear operations to achieve computational power. The issue of non-linearity is well-described already in the reverse correlation literature, and its consequences have been clearly described (Theunissen et al., 2000). There are extensions to the reverse correlation technique to describe lower-order non-linear interactions in the input space (Neri & Heeger, 2002). Such extensions could be applied to the interpretation of classifier’s features. Interestingly, the reverse correlation approach bears some similarities with the “distillation” method from the machine learning literature (Hinton et al., 2015). Distillation consists in mimicking the behavior of a black-box classifier with an easily-interpretable classifier, such as a linear one (linear SVM, etc.). Both techniques can thus be viewed as attempting to find linear approximations of a classifier’s operations, but their precise relationship remain to be investigated.

Second, the method has a number of free parameters: the probability and size of bubbles; the space to generate the probing noise with reduction dimension methods such as PCA when the training set is available. These are hyperparameters that, in our implementation, are not algorithmically constrained. In the examples above, the parameter space was explored heuristically. One suggested heuristic was to try and cover all output classes in a more or less balanced manner with the probing set. We have not developed any fitness criterion, i.e. a way to quantify the efficiency of the method for a given set of parameters. We do provide a quantitative metric based on an insertion/deletion technique (Petsiuk et al., 2018) that could conceivably be used as a fitness criterion if needed. However, we would argue that the iterative process for parameter tuning can be part of the interpretation process since finding the right probing structure in itself provides some information on the structure of the dataset. Also, assessing whether the discovered features make intuitive sense relies mostly on the knowledge and goals of the user.

Third, the method implicitly assumes that there are no invariances by translation in the classifier’s algorithm. With reverse correlation, each point of the input space is treated independently of all others, so a feature discovered in one sub-part of the input space will not impact other, perhaps similar features in other sub-parts. This assumption is obviously falsified by CNN architectures, which are purposely designed to incorporate such invariances. In the CNN example illustrated here with digits recognition, this limitation was circumvented by the fact that all digits in the probing set were roughly spatially aligned. For the SVM on audio data, a time-averaging over the time dimension achieved a similar effect. For the EEG and sleep classification, spectral analysis was used. Thus, mitigation strategies are available: a rough alignment of the probing data (spatially or temporally) should be sufficient for the reverse correlation to produce meaningful results, or a transform to a more robust representation. Another possible direction to address these invariance issues is to generate the probing noise in an appropriate space. Using a pre-processing PCA partly achieves this.

Fourth, here we probed the classifier in the same representation used to train the classifier, even though that representation was not necessarily the raw input representation. This can be extended by probing the classifier in a space other than the training input space and applying an additional transform to feed the train classifier. For instance, as has been suggestion in visual studies using bubbles, spatial frequencies could be sampled with bubbles (e.g., Willenbockel et al., 2010; Caplette et al., 2014), transformed back into pixel images and fed to a classifier trained on images.

### 3.4 Perspectives

The probing method is technically applicable to any classifier’s architecture with any kind of input data. It is thus beyond the scope of this final section to list all possible use cases in the context of neuroscience. We will simply provide a few suggestions, to illustrate perspectives of improvement and the kind of problems that could benefit from the probing method.

In the case of studying perceptual decisions, one possible insight gained from interpreting a classifier is the exploration of the input representation fed to the classifier. The hypothesis is that, the more appropriate the representation, the more explainable the classifier should be. For instance, one could assume that the massively non-linear transformations of auditory and visual information that characterize perceptual systems serve to build a stimulus manifold within which perceptual boundaries are approximately linear (Georgopoulos et al., 1986; Jazayeri & Movshon, 2006; Kell et al., 2018). So, with the correct representation, a classifier modeling a perceptual decision process should be easily interpretable, or at least more easily interpretable than if the input representation was not reflecting perceptual processing. It is with this hypothesis in mind that the audio samples of the example illustrated above were first processed with an auditory model. Even though there are successful deep learning models operating on the raw audio waveforms (e.g. Wavenet, Oord et al., 2006), it is not expected that interpreting them in terms of waveform features will be meaningful. For instance, inaudible phase shifts between frequency components in the input would impact the waveform representation, but should not change the classifier’s decision. An auditory model, in contrast, incorporates transforms inspired by the neurophysiology of the hearing system. If the features extracted resemble those available to a human observer, then they should be revealed when probing a classifier. In fact, the ease of interpreting a classifier feature could be a proxy to evaluate an input representation’s adequation to a perceptual task.

Another possible application concerns the construction of “ideal observer” models (Geisler, 2004). The idea of an ideal observer model is to compute the best theoretical performance on a task, given a set of assumptions (classically, endowing the ideal observer with unbiased decision criteria, perfect and unlimited memory, and so on). This upper performance boundary is then compared to the observed performance with human participants or neural recordings. When considering classification or discrimination tasks, and when a formal model of the ideal observer is unavailable, it can be of interest to build pseudo-ideal observer models with machine learning classifiers. The advantage of our probing method is then that the classifier’s strategy can be directly compared to a reverse correlation analysis of neural or psychophysical data, to ask whether the classifier and the experimental observer used the same decision features.

Finally, the general benefits of interpreting classifiers also apply to the field of neuroscientific applications. In a broad sense, probing is intended to help an expert making sense of a classifier’s strategy. If the features discovered through probing fit a theoretical model, this would reassure the expert that the performance relies on reasonable principles, which is especially important in clinical applications. In return, the expert’s intuition may also help improve the classifier, for instance by simplifying its input representation through pre-processing, and so hopefully making it less brittle to irrelevant variations in input that may have been picked up by overfitting during training (Goodfellow et al., 2015). The discriminative features could be particularly useful to reduce the complexity of a classifier. Based on the discriminative features map, it may be possible to select a subset of important and intelligible features, which can then be used to build a more computationally efficient classifier, for very large dataset and/or for real-time processing.

## 4 Conclusions

We presented a novel method to interpret machine-learning classifiers that is agnostic, versatile and well-suited to applications in the neuroscience domain. Based on the reverse correlation framework, our method uses stochastic perturbation of inputs to observe the classifier’s output. It then visualizes, in the input space, the discriminative and represented features discovered by the classifier for each category. This method can be applied to any kind of classifier. It displays the same well-established benefits and limitations as reverse correlation as applied to psychophysical or neural data. Our hope is that such a method can provide a simple and generic interface between neuroscientists and machine-learning tools.

## Acknowledgements

Research supported by grants ANR-16-CONV-0002 (ILCB), ANR-11-LABX-0036 (BLRI) and the Excellence Initiative of Aix-Marseille University (A*MIDEX) (ET), ANR-10-LABX-0087 IEC and ANR-10-IDEX-0001-02 PSL (DP). TA was supported by the Human Frontier Science Program (LT000362/2018-L).

## Declarations of interest

none.

## Supplementary Materials

**Supplemental Figure 1.**
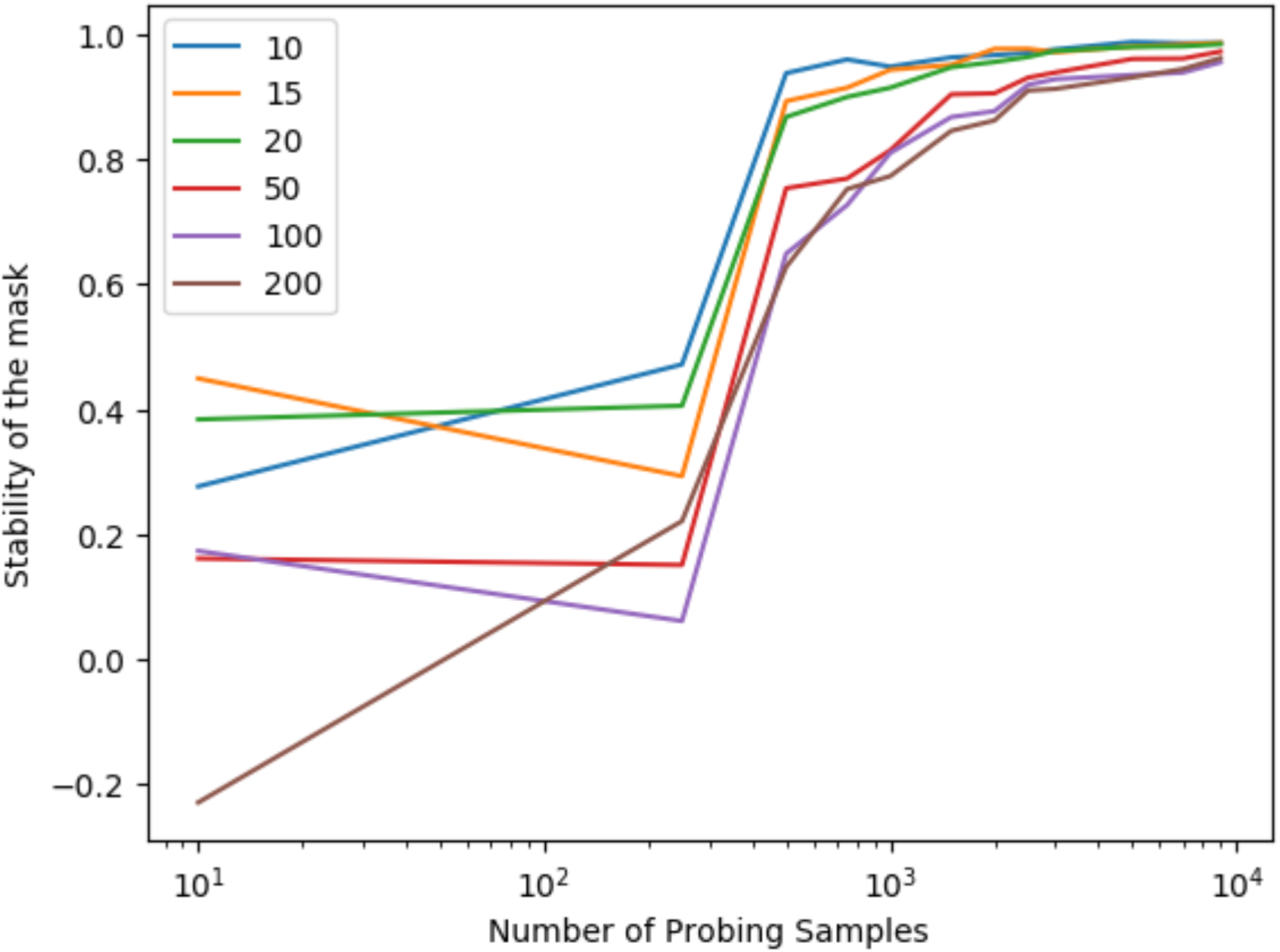
Influence of the number of probing samples on the stability of the mask in the digit classification example (Method 1b, section 2.1.2). The stability corresponds to the correlation between the interpretation maps obtained between two successive number of probing samples. Each curve corresponds to a different number of PCA dimensions used to generate the pseudo random noise. An order of magnitude of 10^3^ to 10^4^ generated probing samples seems sufficient to obtain a stable estimation. It is noticeable that the maps obtained with a smaller number of PCA dimensions converge more quickly than those obtained with more dimensions.

**Supplemental Figure 2.**
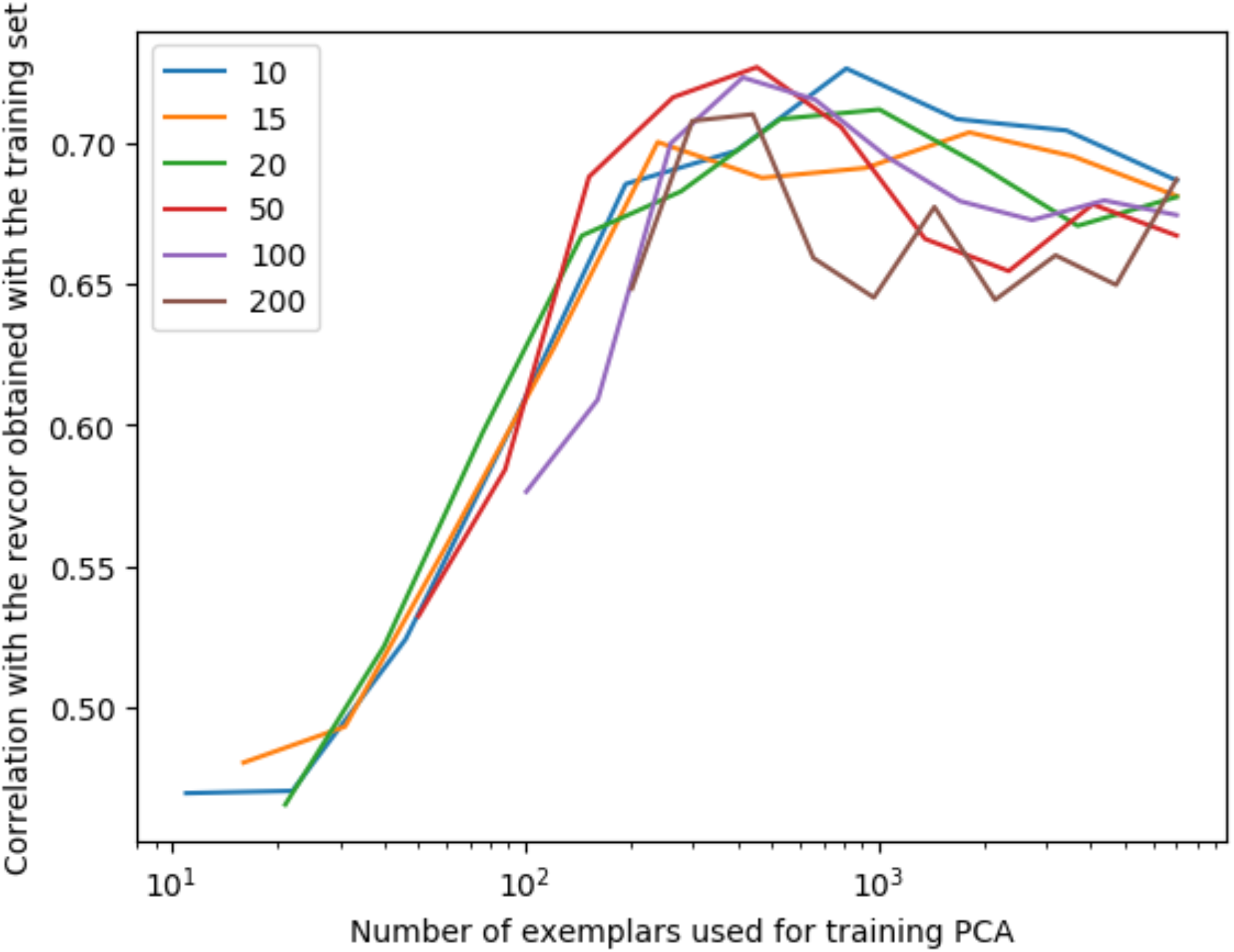
Influence of the number of exemplars used to obtain the PCA for generating the pseudo-random probing samples in the digit classification example (Method 1b, section 2.1.2). Each curve corresponds to a different number of dimensions retained in the PCA. 10^2^ to 10^3^ test samples seem sufficient when the training set is unavailable to converge toward a map correlating with the reference one obtained when the training set is available.

**Supplemental Figure 3.**
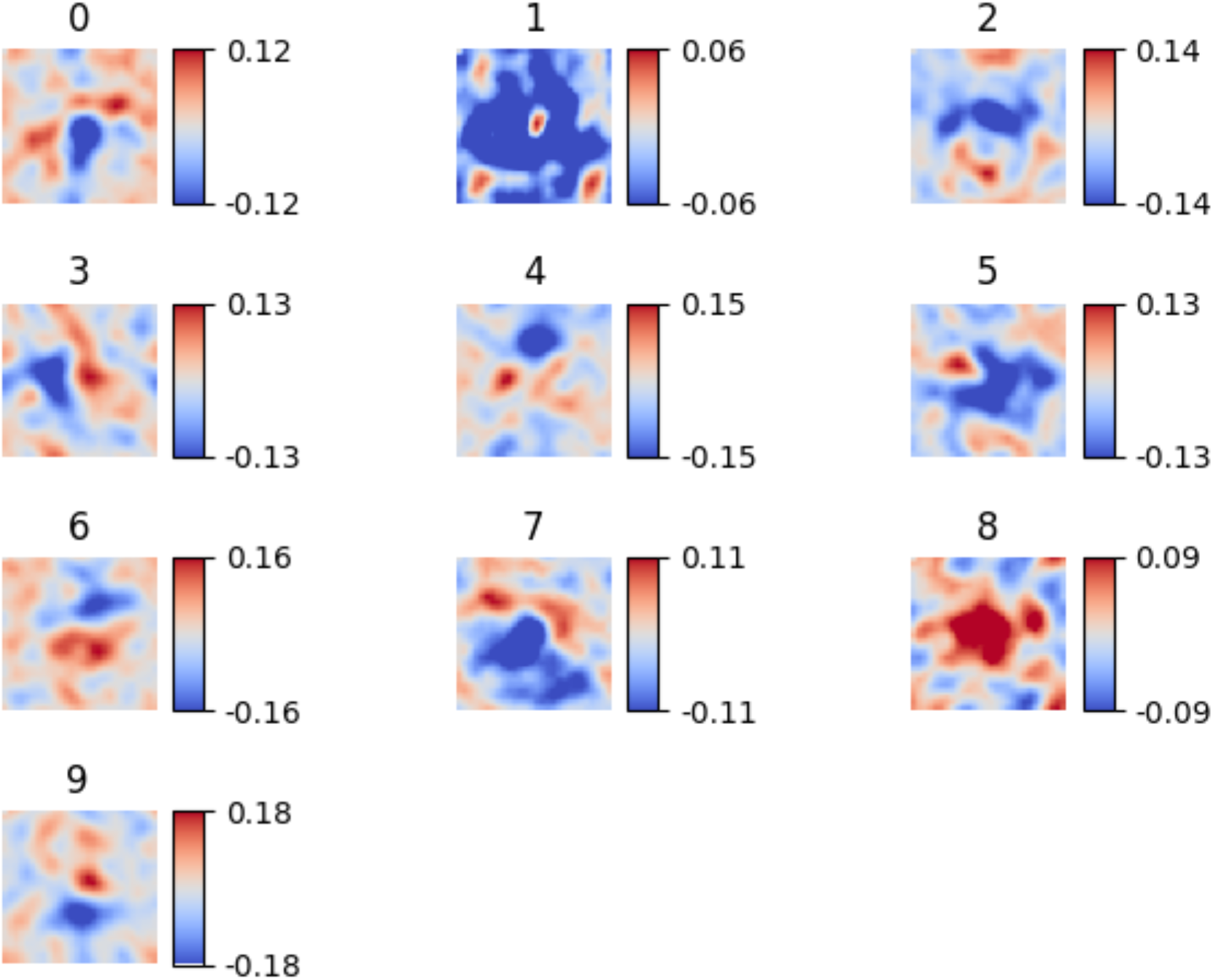
Represented features maps for a CNN classifying handwritten digits for method 2a obtained with a Gabor Noise (Brinkman et al., 2017). Methods 2a and 2b (Figure 4) actually provided correlated maps (average Pearson coefficients for the 10 maps: M = .43 (SD = .10), *df* = 783, all *p* < 10^−3^).

**Supplemental Figure 4.**
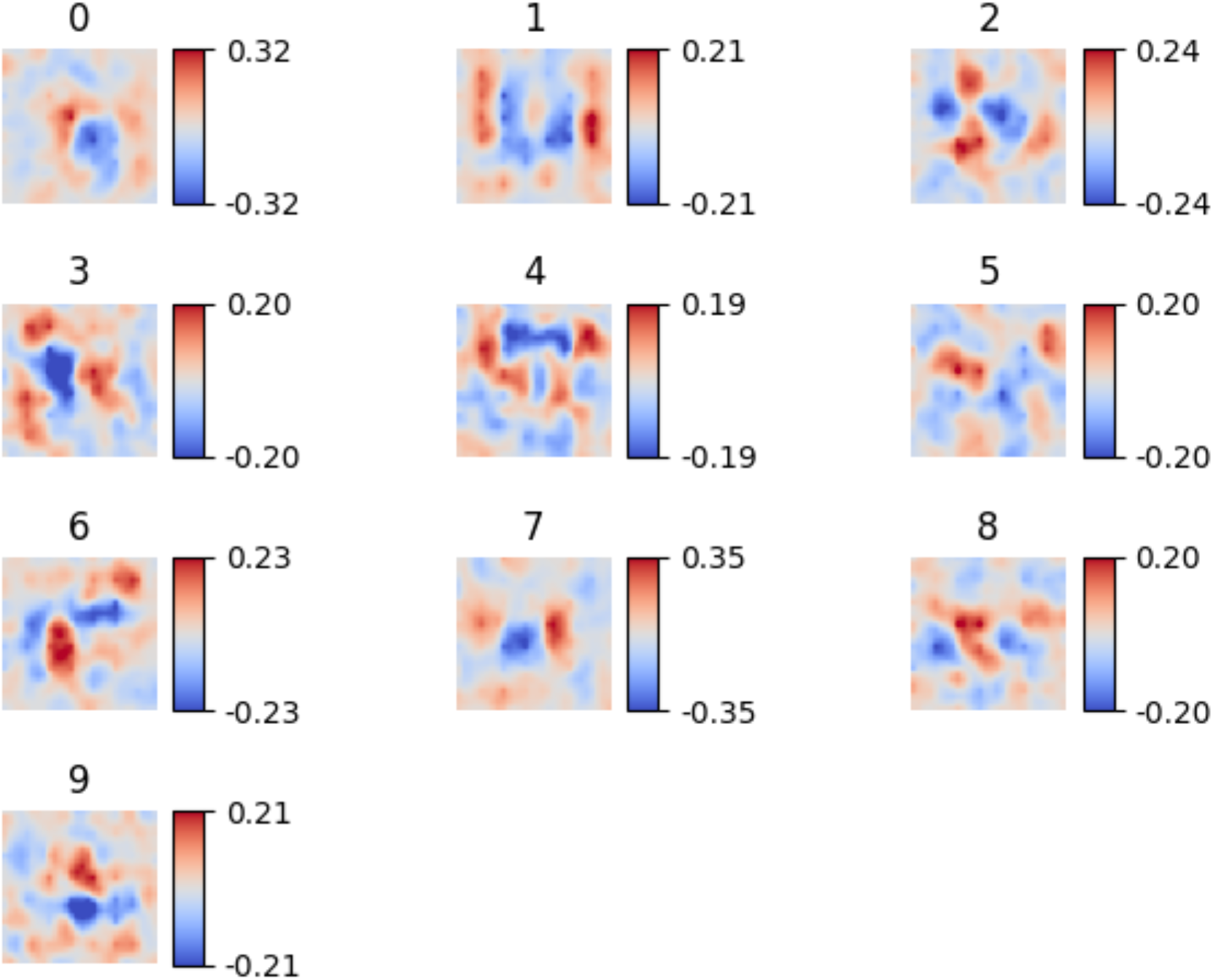
Represented features maps for a CNN classifying handwritten digits for method 2a obtained with a Fractal Noise (Perlin & Hoffert. 1989). Methods 2a and 2b (Figure 4) actually provided correlated maps (average Pearson coefficients for the 10 maps: M = .47 (SD = .08), *df* = 783, all *p* < 10^−3^).

## Footnotes

1 We wish to thank an anonymous reviewer for this suggestion.

2 For clarity here and for illustrating the consequences of the choice of noise, variant a) of the methods will always use Gaussian noise whereas variant b) will always use PCA-based noise, but in practice, other combinations are possible.

## Notes

https://github.com/EtienneTho/proise/

### Competing Interest Statement

The authors have declared no competing interest.

